# Revisiting the role of synaptic plasticity and network dynamics for fast learning in spiking neural networks

**DOI:** 10.1101/2021.01.25.428153

**Authors:** Anand Subramoney, Guillaume Bellec, Franz Scherr, Robert Legenstein, Wolfgang Maass

**Affiliations:** Institute for Theoretical Computer Science, Graz University of Technology, Austria; Institute for Neural Computation, Ruhr University Bochum, Germany; Laboratory of Computational Neuroscience, Ecole Polytechnique Fédérale de Lausanne (EPFL), Switzerland

## Abstract

Spike-based neural network models have so far not been able to reproduce the capability of the brain to learn from very few, often even from just a single example. We show that this deficiency of models disappears if one allows synaptic weights to store priors and other information that optimize the learning process, while using the network state to quickly absorb information from new examples. For that, it suffices to include biologically realistic neurons with spike frequency adaptation in the neural network model, and to optimize the learning process through meta-learning. We demonstrate this on a variety of tasks, including fast learning and deletion of attractors, adaptation of motor control to changes in the body, and solving the Morris water maze task – a paradigm for fast learning of navigation to a new goal.

**Significance Statement:** It has often been conjectured that STDP or other rules for synaptic plasticity can only explain some of the learning capabilities of brains. In particular, learning a new task from few trials is likely to engage additional mechanisms. Results from machine learning show that artificial neural networks can learn from few trials by storing information about them in their network state, rather than encoding them in synaptic weights. But these machine learning methods require neural networks with biologically unrealistic LSTM (Long Short Term Memory) units. We show that biologically quite realistic models for neural networks of the brain can exhibit similar capabilities. In particular, these networks are able to store priors that enable learning from very few examples.

## 1 Introduction

Modelling and theoretical investigation of learning capabilities of models for neural networks of the brain, in particular of networks of spiking neurons, has focused on learning via synaptic plasticity, such as spike-timing-dependent plasticity (STDP). But experimental data suggest that synaptic plasticity does not capture all learning capabilities of neural networks in the brain. In particular, brains can learn very fast, even in a single trial (Brea and Gerstner, 2016), whereas STDP requires numerous repetitions of a trial (Froemke et al., 2010). It has also been argued, see for example the comments to Suppl. Figure 3 in (Wang et al., 2018), that there is no direct evidence that dopamine can drive synaptic change this rapidly for fast reinforcement learning. On the other hand, a number of experimental data suggest that brains use, in addition to or instead of synaptic plasticity, the dynamics of network states to store new information (Perich et al., 2018; Botvinick et al., 2019; Crevecoeur et al., 2020a).

But it has remained open how data-driven models for neural networks of the brain could achieve this. In the meanwhile, a functionally powerful form of fast learning without synaptic plasticity has been demonstrated in particular types of artificial neural network: in networks of Long Short-Term memory (LSTM) units (Hochreiter et al., 2001; Duan et al., 2016; Wang et al., 2016, 2018). Unfortunately both the mechanism of LSTM units and their activity patterns cannot be directly related to biophysical mechanisms or recorded activity patterns of neural networks in the brain. In particular, LSTM units employ registers, similar to digital computers, for rapidly storing information for an indefinite amount of time. Furthermore, an LSTM unit only needs to become active when new information is stored, updated, or read out.

However recently it has been shown that a substantial fraction of the functional capability of networks of LSTM units can be reproduced by networks of spiking neurons, provided that they also contain neurons with spike frequency adaptation (SFA) (Bellec et al., 2018b; Salaj et al., 2020). SFA means that a neuron increases its firing threshold after firing. Experimental data from the Allen Institute (Allen Institute, 2018) show that a substantial fraction of excitatory neurons of the neocortex, ranging from 20% in mouse visual cortex to 40% in the human frontal lobe, exhibit SFA. We show that neurons with SFA endow networks of spiking neurons with the capability to learn very fast, even without synaptic plasticity. We focus on two characteristic aspects of the resulting new learning theory for networks of spiking neurons:

1. Synaptic weights are able to encode instructions for controlling learning processes, in particular learning processes that do not rely on synaptic plasticity. This perspective enables brains to employ a much larger and functionally more powerful repertoire of learning schemes, going far beyond what can be learnt by just using synaptic plasticity.
2. Synaptic weights are also able to encode priors, in particular priors that enable efficient learning from few examples by exploiting common structure in families of related learning tasks (Gershman and Niv, 2010).

We demonstrate each of these two principles separately in two illustrative tasks (see Fig. 2 and 4) and together in applications to standard motor control and navigation tasks that require self-supervised learning and reinforcement learning (Fig. 3 and 5).

In line with (Harlow, 1949; Wang et al., 2016, 2018; Zador, 2019) we focus on a setting where synaptic weights are tuned on a large time-scale that conceptually reflects evolutionary and developmental processes as well as prior learning. We show that this setup can also be used to elucidate fast learning capabilities of spiking neural networks, i.e., of biologically more realistic models for neural networks of the brain.

## 2 Methods

### 2.1 Network models

Neurons were modelled after the standard leaky integrate-and-fire (LIF) model with a proportion of neurons in all the networks consisting of LIF neurons with spike frequency adaptation (SFA) as in (Bellec et al., 2018b; Salaj et al., 2020) and described here. **Leaky integrate and fire (LIF) neurons**. A LIF neuron *j* has one state variable – its membrane potential *V*_*j*_(*t*). The neuron spiked whenever the membrane potential *V*_*j*_(*t*) exceeded the threshold *v*_th_. At each spike time *t*, the membrane potential *V*_*j*_(*t*) was reset by subtracting the threshold value *v*_th_. After this, the neuron entered a refractory period of some *τ*_ref_ time steps during which time it cannot spike. Between spikes, the membrane potential *V*_*j*_(*t*) evolved according to:

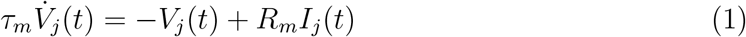

where *I*_*j*_(*t*) is the input, and *R*_*m*_ is the electrical resistance term.

In discrete time, the input and output spike trains were modeled as binary sequences *x*_*i*_(*t*), *z*_*j*_(*t*) ∈ {0, 1}. Neuron *j* emitted a spike at time *t* if it was currently not in a refractory period, and its membrane potential *V*_*j*_(*t*) was above its threshold. During the refractory period following a spike, *z*_*j*_(*t*) was fixed to 0. In discrete time, using timesteps of *δt*, the neuron was simulated as:

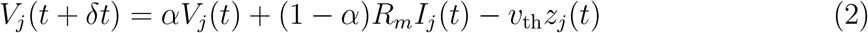

where 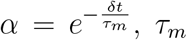, *τ*_*m*_ is the membrane constant of the neuron *j*. The neuron spike is defined as *z*_*j*_(*t*) = *H*(*V*_*j*_(*t*) − *v*_th_), where *H*(*x*) is the Heaviside step function i.e. *H*(*x*) = 1 if *x >* 0 and 0 otherwise. In all our simulations, *δt* was set to 1 ms and *R*_*m*_ was set to 1 GΩ.

The input current *I*_*j*_(*t*) was defined as the weighted sum of spikes from external inputs and other neurons in the network:

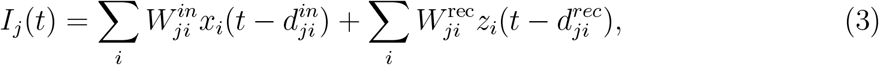

where 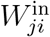 and 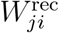 denote respectively the input and the recurrent synaptic weights and 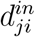 and 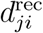 the corresponding synaptic delays from neuron *j* to neuron *i*. **LIF neurons with SFA**. A SFA mechanism was added to the LIF neuron model in accordance with the *GLIF*_2_ neuron model of (Teeter et al., 2018) by replacing the fixed threshold *v*_th_ with an adaptive threshold *A*_*j*_(*t*). Whenever the membrane potential *V*_*j*_(*t*) exceeded the threshold *A*_*j*_(*t*) (instead of *v*_th_), this neuron emitted a spike *z*_*j*_, and its membrane potential was reset as before.

*A*_*j*_(*t*) followed the dynamic:

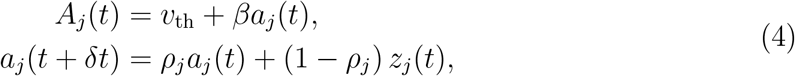

where *a*(*t*) is its activity-dependent component, *β >* 0 is the relative amplitude of the activity-dependent component. Essentially, after each spike, the adaptive threshold *A*_*j*_(*t*) contains a variable component *a*_*j*_(*t*) that is increased by a constant and is allowed to decay back to 0. The parameter 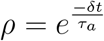 controls the speed by which *a*(*t*) decays back to 0, where *τ*_*a*_ is the adaptation time constant. Adaptation time constants of neurons with SFA were chosen to match the task requirements

### 2.2 Details of the learning to learn setup

#### Network simulation in the inner loop

In each episode of the inner loop, a task *C* was chosen from the family of tasks ℱ. The RSNN received a sequence of inputs x^*k*^ corresponding to this *C*, each encoded through the population activity of spiking neurons. In addition, it received either a cue (experiment in Section 3.2) or feedback (all other experiments) of what the target output should have been for the previously presented input *C*(*x*^*k−*1^) (The feedback was set to zero in the first time step). The network could use the cue or delayed target feedback to adapt its behavior. The network had to predict the target *y*^*k*^ = *C*(*x*^*k*^) at each time step *k*. There was no synaptic plasticity in the inner loop.

#### Hyperparameter optimization in the outer loop

The outer loop optimization of learning-to-learn happens in the following way: In each iteration, a batch of different random tasks are chosen from the family ℱ and the corresponding inputs are presented to the RSNN in a inner loop. The predictions from the inner loop are used to compute a loss function that compares the prediction to the target, for the entire batch of tasks. We used back-propagation through time (BPTT) to optimize the hyperparameters in the outer loop of L2L, which are the synaptic weights of the RSNN in our applications.

Since the spike output of a LIF neuron model is not differentiable, we used a pseudo-derivative with an additional factor *γ <* 1 that dampens the increase of backpropagated errors through spikes as in (Bellec et al., 2018b; Salaj et al., 2020):

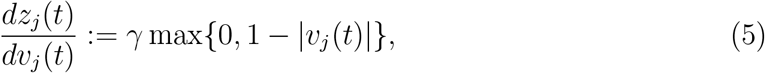

where *v*_*j*_(*t*) denotes the normalized membrane potential 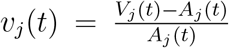. A proper choice of the dampening factor turns out to be critical in such applications of BPTT to RSNNs, since the gradient needs to propagate backwards through many layers (= time slices) of the unrolled RSNN. In neurons with SFA, gradients can be propagated efficiently through the hidden variable that denotes the dynamic threshold, without requiring a pseudo-derivative or dampening factor like for the backpropagation through spikes.

### 2.3 Details of the learning experiments in Results

#### 2.3.1 New learning capabilities of recurrent networks of spiking neurons

##### Task family

In each task *C* from the family of tasks ℱ, three 25-bit randomly generated patterns were shown in the first phase referred to as phase A (see Fig. 2 for illustration of phases). Each of these three patterns were presented to the network for 100 ms. In the next phase B, partial versions of the same three patterns were shown. The partial patterns were generated by setting each non-zero bit in each pattern to zero with 40% probability. The targets for this phase were the full patterns from phase A. In phase C, a cue signal was given to denote to the network which of the three patterns should be “deleted” by index. In the last phase D, the same partial patterns as in phase B were shown again. The targets were the full patterns for the two patterns that were not “deleted”. For the pattern that was “deleted”, the target was one of other two patterns that was closest to the deleted pattern measured by hamming distance.

##### Input encoding

The patterns were generated as 25-bit vectors, and the cue as a 3-bit vector. In both cases, each bit was represented by 5 spiking neurons, which fired at a high rate of 200 Hz when the bit was 1, and at a lower rate of 2 Hz when it was 0.

##### Output decoding

The output of the RSNN was a linear readout that received as input the trace of the firing of all the neurons in the network. The spiking activity of the neurons was convolved with an exponential kernel with time constant 100 ms to generate the trace.

##### RSNN setup and training schedule

The standard RSNN architecture was used, with 300 hidden neurons. Of these, 50% were LIF neurons with SFA and the rest were LIF neurons without SFA. We used all-to-all connectivity between all neurons.

The network training proceeded as follows: In each episode, a new set of three random patterns were chosen along with the pattern that was to be excluded from the output. These were presented to the network as described above.

All the weights of the RSNN were updated using our variant of *BPTT*, once per iteration, where an iteration consists of a batch of 100 episodes, and the weight updates were accumulated across episodes in an iteration. We used the Adam optimizer (Kingma and Ba, 2014) with the default parameters with a learning rate of 0.001. The loss function for training was the bit-wise cross-entropy loss of the RSNN predictions:

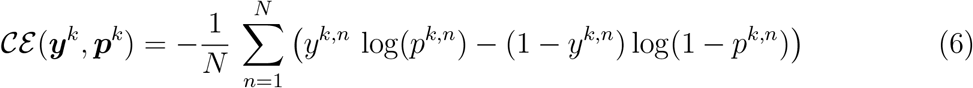

where 𝒞*ε* is the cross-entropy loss, ***y***^*k*^ and ***p***^*k*^ are, respectively, the target 25-bit vector and vector of predicted probability of each of the *N* = 25 bits being 1 at step *k*.

The overall loss was given by:

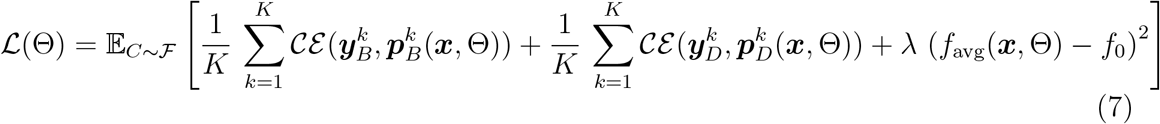

where *K* = 3 is the number of patterns shown in each phase, and the subscripts *B* and *D* denote phase to which the target and prediction vectors correspond, *λ* = 5 is the coefficient of the regularization term, *f*_avg_ is the average firing rate of the network over the entire episode, and *f*_0_ = 20 Hz is the target firing rate in the regularization term. With the regularization term, we induce the RSNN to use sparse firing. We trained the RSNN for 100,000 iterations.

##### Parameter values

The RSNN parameters were as follows: 5 ms neuronal refractory period, delays of 1 ms, adaptation time constants of the LIF neurons with SFA were spread uniformly between 1 − 1000 ms, *β* =1.7 mV for LIF neurons with SFA (0 for LIF neurons without SFA), membrane time constant *τ* = 20 ms, 30 mV baseline threshold voltage. The dampening factor for training was *γ* = 0.3.

#### 2.3.2 Fast adaptation of motor predictions

##### Task family

The family of functions was defined by different two-link arms where the length and masses of the links were randomly chosen in the range [0.5, 2]. The torques were generated randomly as described in (Gilra and Gerstner, 2017). The network was trained to predict the arm state, i.e. the angles *ϕ*_1_, *ϕ*_2_ of its two links.

##### Input encoding

Analog values were transformed into spike trains to serve as inputs to the RSNN as follows: For each input component, 100 input neurons are assigned values *c*_1_, …, *c*_100_ evenly distributed between the minimum and maximum possible value of the input. Each input neuron has a Gaussian response field with a particular mean and standard deviation, where the means are uniformly distributed between the minimum and maximum values to be encoded, and with a constant standard deviation. More precisely, the firing rate ri (in Hz) of each input neuron i is given by 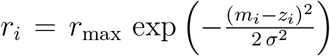, where *r*_max_ = 200 Hz, *m*_*i*_ is the value assigned to that neuron, *z*_*i*_ is the analog value to be encoded, and 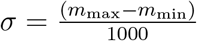, *m*_min_ with *m*_max_ being the minimum and maximum values to be encoded.

##### Output decoding

The output of the RSNN was a linear readout that received as input the trace of the firing of all the neurons in the network. The spiking activity of the neurons was convolved with an exponential kernel with time constant 50 ms to generate this trace. *RSNN setup and training schedule:* The standard RSNN model was used, with 600 hidden neurons. Of these, 50% were LIF neurons with SFA and the rest were LIF neurons without SFA. We used all-to-all connectivity between all neurons.

The training was as follows: During inner loop training, for each episode, we randomly chose a value for the mass and length for each link of the arm. The RSNN received the motor command *c*(*t*) = [*c*_1_(*t*) *c*_2_(*t*)]^*T*^, and the actual state vector of the arm *τ* = 100ms ago, *s*(*t*−*τ*) as inputs. The state vector of the arm *s*(*t*) = [*ϕ*_1_(*t*) *ϕ*_2_(*t*)]^*T*^ was defined by the angles *ϕ*_1_, *ϕ*_2_ of its two links. All the inputs were encoded into spikes using a population-rate code before being presented to the network (as shown in Fig. 3D top panel). A linear readout on the trace of the neural activity was used to generate the predictions of the state of the arm *s*^(*t*; Θ). Each episode lasted for 30 seconds, where the torque changed every 10 ms.

In the outerloop, the following loss function was minimized using *BPTT* for spiking networks:

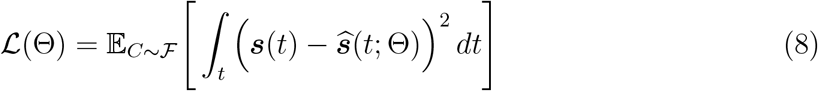

##### Parameter values

The RSNN parameters were as follows: 5 ms neuronal refractory period, delays of 1 ms, adaptation time constants of the LIF neurons with SFA spread uniformly between 1 −600 ms, *β* = 1.7 mV for LIF neurons with SFA (0 for LIF neurons without SFA), membrane time constant *τ* = 20 ms, 30 mV baseline threshold voltage. The dampening factor for training was *γ* = 0.3. We used the Adam optimizer (Kingma and Ba, 2014) with the default parameters with a learning rate of 0.001 and a batch size of 80 for training.

#### 2.3.3 Priors encoded in synaptic weights can significantly speed up learning

##### Task family

RSNN was trained to implement a regression algorithm on a family of sinusoidal functions. The targets were defined by sinusoidal functions *y* = *A* sin(*ϕ* + *x*) over the domain *x* ∈ [− 5, 5]. The specific function to be learned was defined then by the phase *ϕ* and the amplitude *A*, which were chosen uniformly random between [0, *π*] and [0.1, 5] respectively.

##### Input encoding

Analog values were transformed into spiking trains in exactly the same way as for the previous section.

##### Output decoding

The output of the RSNN was a linear readout that received as input the mean firing rate of each of the neurons per step i.e the number of spikes divided by 20 for the 20 ms time window that constitutes the step.

##### RSNN setup and training schedule

The standard RSNN model was used, with 100 hidden neurons, of which 40% were LIF neurons with SFA and the rest were LIF neurons without SFA. We used all-to-all connectivity between all neurons.

##### The network training proceeded as follows

A new target function was randomly chosen for each episode of training, i.e., the parameters of the target function were chosen uniformly randomly from within the ranges above. Each episode consisted of a sequence of 500 steps, each lasting for 20 ms. In each step, one training example from the current function to be learned was presented to the RSNN. In such a step *k*, the inputs to the RSNN consisted of a randomly chosen scalar input *x*^*k*^. In addition, at each step, the RSNN also got the target value *C*(*x*^*k−*1^) from the previous step, i.e., the value of the target calculated using the target function for the inputs given at the previous step (in the first step, *C*(*x*^0^) is set to 0).

All the weights of the RSNN were updated using our variant of *BPTT*, once per iteration, where an iteration consists of a batch of 100 episodes, and the weight updates were accumulated across episodes in an iteration. We used the Adam optimizer (Kingma and Ba, 2014) with the default parameters with a learning rate of 0.001. The loss function for training was the mean squared error (MSE) of the RSNN predictions over an iteration (i.e. over all the steps in an episode, and over the entire batch of episodes in an iteration):

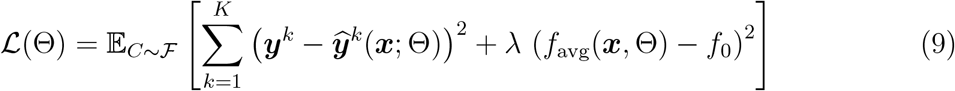

where *K* = 500 is the number of steps i.e. the number of points presented to the network, each lasting for 20 ms, 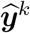 and ***y***^*k*^ are the network prediction and target respectively at each step *k*, ***x*** is the input to the network, *λ* = 30 is the coefficient of the regularization term, *f*_avg_ is the average firing rate of the network over the entire episode, and *f*_0_ = 20 Hz is the target firing rate in the regularization term. With the regularization term, we induce the RSNN to use sparse firing. We trained the RSNN for 5000 iterations.

##### Parameter values

The RSNN parameters were as follows: 5 ms neuronal refractory period, delays of 1 ms, adaptation time constants of the LIF neurons with SFA spread uniformly between 1− 3000 ms, *β* = 1.6 mV for LIF neurons with SFA (0 for LIF neurons without SFA), membrane time constant *τ* = 20 ms, 30 mV baseline threshold voltage. The dampening factor for training was *γ* = 0.3.

##### Analysis and comparison

The linear baseline was calculated by performing linear regression on the analog values of input points and targets in the first half of the episodes (250 steps) and testing it on the points in the second half of the episode. The total test MSE was 0.1968 ±0.1469 and the linear baseline was 4.0340.

For visualizing the internal model of the RSNN, we show, for any potential input value *x*, the output *y* which the RSNN would give if this *x* would occur as the next network input (in a hypothetical experiment, that has no effect on the next steps of the learning process for learning the target function *f*). More precisely, to produce these panels, we stored the network state (i.e., the membrane potentials and all other dynamic parameters) at the corresponding time steps during the inner-loop learning process. We then continued the simulation from these states with inputs from -5 to 5 and the network predictions were plotted as the orange curve in in panels Fig. 4B-E. The network state was not allowed to change when these test inputs *x* are shown.

#### 2.3.4 Spiking neural networks can learn extremely fast from rewards — without engaging synaptic plasticity

##### Task family

The tasks consisted of a family of navigation tasks in a two-dimensional circular arena. For all tasks, the arena was a circle with radius 1 and goals were smaller circles of radius 0.3 with centers uniformly distributed on the circle of radius 0.85. At the beginning of an episode and after the agent reached a goal, the agent’s position was set randomly with uniform probability within the arena. At every timestep, the agent chose an action by generating a small velocity vector of Euclidean norm smaller or equal to *a*_scale_ = 0.02. When the agent reached the goal, it received a reward of 1. If the agent attempted to move outside the arena, it received a negative reward of −0.02 and its new position was given by the intersection of the velocity vector with the border.

##### Input encoding

Information of the current environmental state *s*(*t*) and the reward *r*(*t*) were provided to the RSNN at each time step *t* as follows: The state *s*(*t*) was given by the *x* and *y* coordinate of the agent’s position (see top of Fig. 5C). Each position coordinate *ξ*(*t*) ∈ [−1, 1] was encoded by 40 neurons which spiked according to a Gaussian population rate code defined as follows: a preferred coordinate value *ξ*_*i*_, was assigned to each of the 40 neurons, where *ξ*_*i*_’s were evenly spaced between −1 and 1. The firing rate of neuron *i* was then given by *r*_max_ exp(−100(*ξ*_*i*_ −*ξ*)^2^) where *r*_max_ was 500 Hz. The instantaneous reward *r*(*t*) was encoded by two groups of 40 neurons (see green row at the top of Fig. 5C). All neurons in the first group spiked in synchrony each time a reward of 1 was received (i.e., the goal was reached), and the second group spiked when a reward of −0.02 was received (i.e., the agent moved into a wall).

##### Output decoding

The output of the RSNN was provided by five readout neurons. Their membrane potentials *y*_*i*_(*t*) defined the outputs of the RSNN. The action vector **a**(*t*) = (*a*_*x*_(*t*), *a*_*y*_(*t*))^*T*^ was sampled from the distribution *π*_*θ*_ which depended on the network parameters *θ* through the readouts *y*_*i*_(*t*) as follows: The coordinate *a*_*x*_(*t*) (*a*_*y*_(*t*)) was sampled from a Gaussian distribution with mean *µ*_*x*_ = tanh(*y*_1_(*t*)) (*µ*_*y*_ = tanh(*y*_2_(*t*))) and variance *ϕ*_*x*_ = *σ*(*y*_3_(*t*)) (*ϕ*_*y*_ = *σ*(*y*_4_(*t*))). The velocity vector that updated the agent’s position was then defined as *a*_*scale*_ **a**(*t*). If this velocity had a norm larger than *a*_*scale*_, it was clipped to a norm of *a*_*scale*_.

The last readout output *y*_5_(*t*) was used to predict the value function *V*_*θ*_(*t*). It estimated the expected discounted sum of future rewards 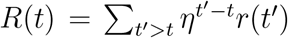, where *η* = 0.99 is the discount factor and *r*(*t*′) denotes the reward at time *t*′. To enable the network to learn complex forms of exploration we introduced current noise in the neuron model in this task. At each time step, we added a small Gaussian noise with mean 0 and standard deviation 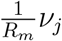 to the current *I*_*j*_ into neuron *j*. Here, *ν*_*j*_ is a network parameter initialized at 0.03 and optimized by *BPTT* alongside the network weights.

##### RSNN setup and training schedule

To train the network we used the Proximal Policy Optimization algorithm (PPO) (Schulman et al., 2017). For each training iteration, *K* full episodes of *T* timesteps were generated with fixed parameters *θ*_*old*_ (here *K* = 10 and *T* = 2000). We write the clipped surrogate objective of PPO as *O*^*PPO*^(*θ*_*old*_, *θ, t, k*) (this is defined under the notation *L*^*CLIP*^ in (Schulman et al., 2017)). The loss with respect to *θ* was then defined as follows:

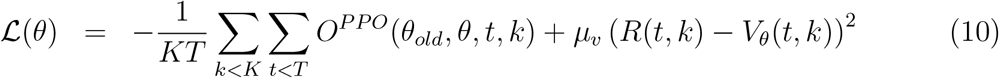

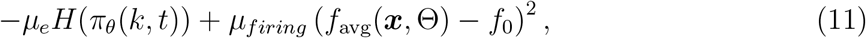

where *H*(*π*_*θ*_) is the entropy of the distribution *π*_*θ*_, *f* ^0^ is a target firing rate of 10 Hz, *µ*_*v*_, *µ*_*e*_, *µ*_*firing*_ are regularization hyper-parameters, *f*_avg_ is the average firing rate of the network over the entire episode. Importantly probability distributions used in the definition of the loss ℒ (i.e. the trajectories) were conditioned on the current noises, so that for the same noise and infinitely small parameter change from *θ*_*old*_ to *θ* the trajectories and the spike trains were the same. At each iteration this loss function ℒwas then minimized with one step of the ADAM optimizer (Kingma and Ba, 2014).

##### Parameter values

In this task the RSNN had 400 hidden units (200 excitatory LIF neurons, 80 inhibitory LIF neurons and 120 LIF neurons with SFA with adaptation time constants *τ*_*a*_ = 1200 ms) and the network was rewired with a fixed global connectivity of 20% using DEEP R (Bellec et al., 2018a). DEEP R provides a way to train a sparsely connected neural network directly using BPPT, while maintaining a fixed overall sparsity in the network and fixed sign for each of the connections. The latter ability allows us to train a network that obey Dale’s law with fixed excitatory and inhibitory neurons. The membrane time constants were similarly sampled between 15 and 30 ms. The adaptation amplitude *β* was set to 1.7. The refractory period was set to 3 ms and delays were sampled uniformly between 1 and 10 ms. The regularization parameters *µ*_*v*_, *µ*_*e*_ and *µ*_*firing*_ were respectively 1, 0.001, and 100. The parameter *E* of the PPO algorithm was set to 0.2. The learning rate was initialized to 0.01 and decayed by a factor 0.5 every 5000 iterations. We used the default parameters for ADAM, except for the parameter *E* which we set to 10^*−*5^.

## 3 Results

We consider recurrent networks of spiking neurons (RSNNs) that contain, besides standard leaky integrate-and-fire (LIF) neurons, also a random subset of neurons with SFA. The network setup across experiments is schematically shown in Fig. 1B. In addition to the input, the network receives either a cue (in the experiment in Sec. 3.2) or a feedback signal (in all other experiments). The network was optimized using learning-to-learning all cases.

**Figure 1:**
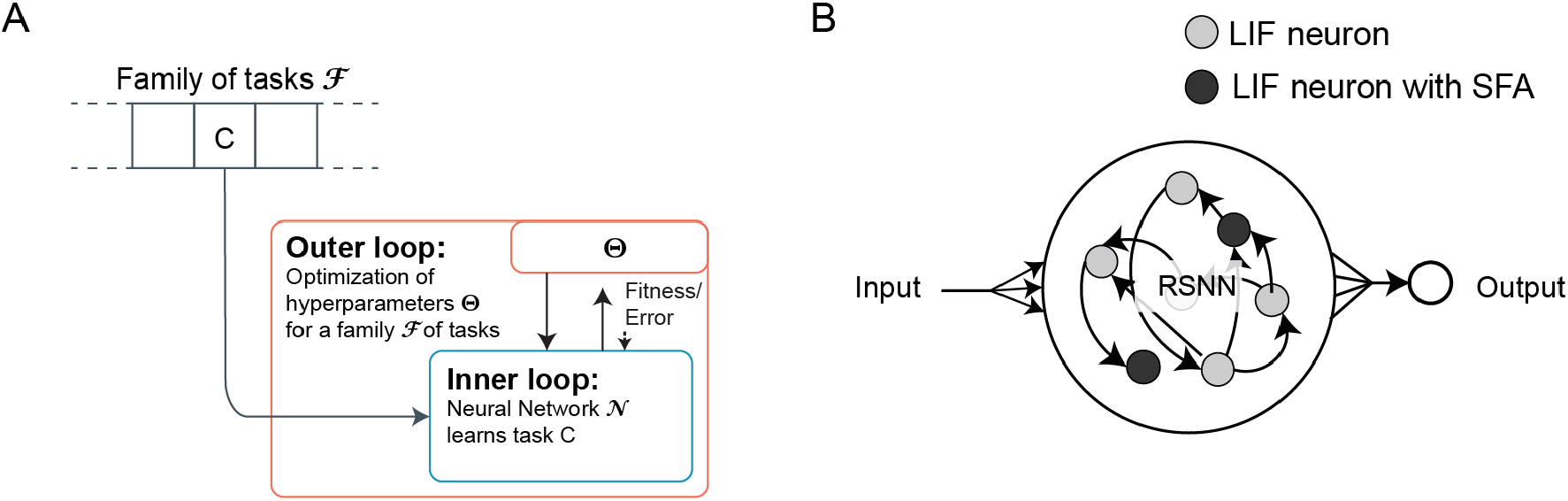
Schema of the learning architecture that we consider. (**A**) The two levels of optimization/learning for learning-to-learn that is used in the experiments in this paper is illustrated here. Learning by a neural network 𝒩 is enhanced by prior optimization of hyperparameters for a large family of learning tasks. (**B**) Generic architecture of the biologically realistic neural network models 𝒩 that we consider in all the experiments, consisting of LIF neurons both with and without SFA.

Experimental data show that SFA produces a history dependence of neural firing on the time scale of seconds, in fact, up to 20 seconds according to (Pozzorini et al., 2013, 2015). The biophysical mechanisms behind SFA include inactivation of depolarizing currents and the activity-dependent activation of slow hyperpolarizing or shunting currents (Gutkin and Zeldenrust, 2014; Benda and Herz, 2003). SFA has already been implicated in cellular short-term memory (Marder et al., 1996; Turrigiano et al., 1996) and other important features of brain networks (Gutkin and Zeldenrust, 2014). We model neurons with SFA as generalized leaky integrate-and-fire neurons, more precisely *GLIF*_2_ neuron models according to (Teeter et al., 2018). Practical advantages of this simple model is that it can be efficiently simulated and that it allows parameter optimization in a neural network model through gradient descent. It is based on the following augmentation of the standard leaky-integrate-and-fire (LIF) neuron model: One adds to the firing threshold *A*(*t*) of a history-dependent component *a*(*t*) that increases by a fixed amount after each spike *z*(*t*) of the neuron, and then decays exponentially back to 0. This history-dependent component models the inactivation of voltage-dependent sodium channels in a qualitative manner.

We first describe the learning-to-learn setup conceptually and then describe the experiments we performed.

### 3.1 Learning to learn

Our analysis builds on a key insight from neuroscience and cognitive science: Fast learning capability of brains is supported by the fact that brains do not learn a new task starting in a tabula-rasa state. Rather, they build on neural circuits, learning skills, and prior knowledge that have been formed throughout evolution, development, and prior learning experiences (Harlow, 1949; Wang et al., 2018; Zador, 2019). The capabilities of this prior shaping of neural circuits can be analyzed with the help of the formal learning-to-learn (L2L) model from (Hochreiter et al., 2001; Wang et al., 2016; Duan et al., 2016) shown in (see Fig. 1A). Optimization (learning) is carried out in this model at two levels: The “inner loop” involves the learning of a single task by a network 𝒩, which will be, in our case, a network of spiking neurons as depicted in Fig. 1B. The “outer loop” involves optimization of some hyperparameters Θ of the network to support fast learning of the individual tasks in the inner loop. The outer loop training proceeds on a much larger time scale than the inner loop, and considers a large, in general infinitely large, range of learning tasks ℱ instead of a single learning task. This outer loop mimics the impact of evolutionary, developmental and prior learning processes, as well as prior learning, on parameters of the neural network 𝒩. Notably, it does not optimize these parameters for a single learning task, but for fast learning of any generic new task *C* from the considered range ℱ of learning tasks. This optimization is carried out in this study through backpropagation through time (BPTT) to minimize the loss on batches of different tasks chosen from the given family of learning tasks ℱ. Note that we use the terms training and optimization interchangeably in this paper. For simplicity, we let all synaptic weights of the RSNN 𝒩 belong to the set of hyperparameters that are optimized in the outer loop. Hence the outer loop training shapes the activation dynamics of the RSNN, which include its firing activity and short-term memory.

### 3.2 New learning capabilities of recurrent networks of spiking neurons

Here, we want to demonstrate point 1 of the Introduction, the substantially enlarged range of learning strategies that become available if one integrates dynamic network states into the learning process. We demonstrate this on some of the arguably most important learning goals for recurrent neural networks: learning an attractor, using a learnt attractor for input completion, and deleting an attractor for pattern completion. The first two learning goals can be achieved through Hebbian learning rules in suitable artificial neural networks such as Hopfield networks (Hopfield, 1982). However the learning of a new attractor typically requires a substantial number of trials, whereas the brain is able to learn a new rule or prototype for image classification in one or very few trials. Deleting an attractor for pattern completion corresponds to learning that a specific rule or prototype is no longer valid. This can also be accomplished by the human brain in one or few trials, but it is difficult to achieve through training of any type of recurrent neural network. But importantly, none of the three mentioned learning goals have been demonstrated for more realistic models of neural networks such as recurrent networks of spiking neurons. We show here that they can achieve all three learning goals very fast, even in a single trial, if one takes into account that synaptic weights can encode a much wider repertoire of learning methods than those that are accessible through local rules for synaptic plasticity such as Hebbian rules or STDP. This is demonstrated in Fig. 2.

**Figure 2:**
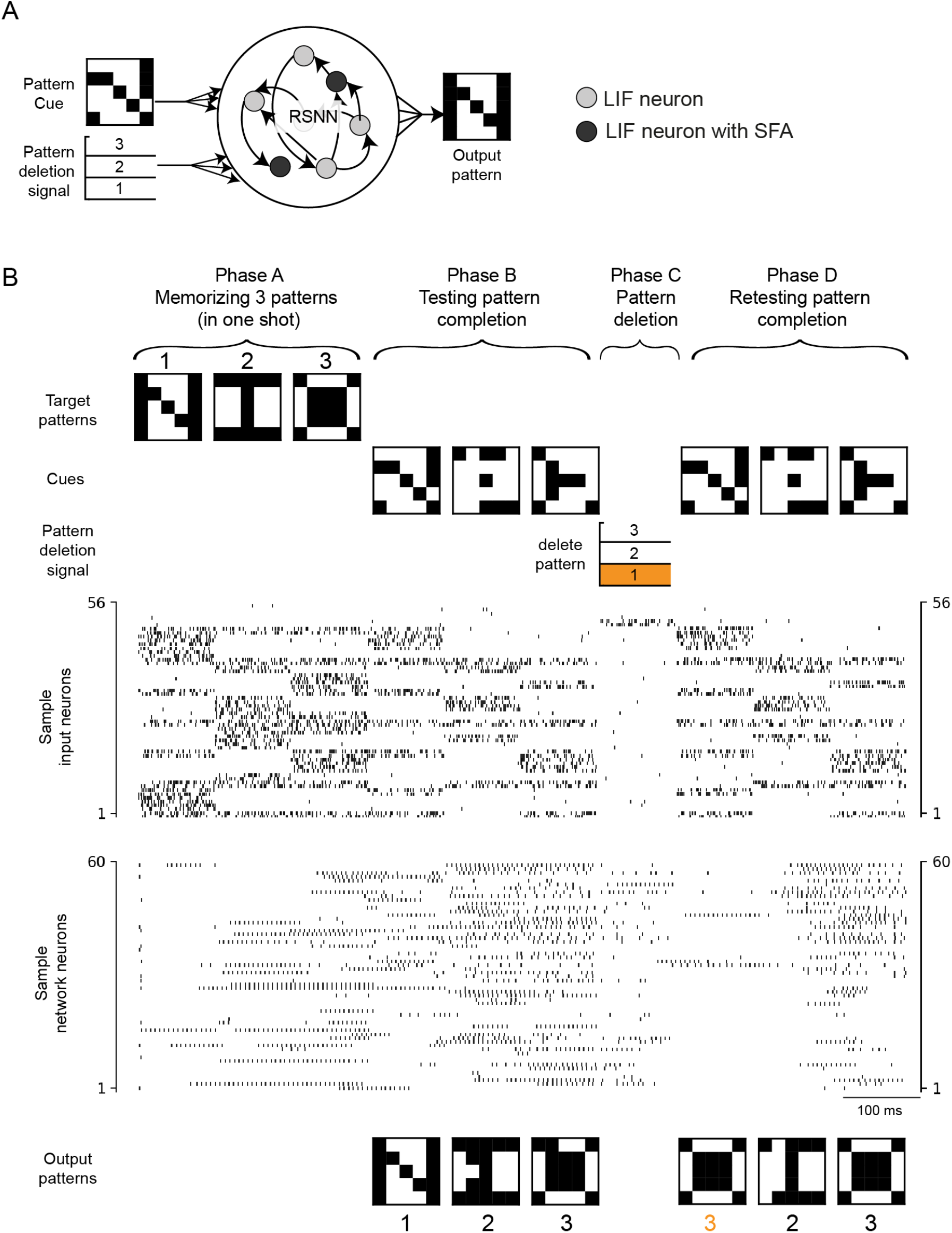
Example of new neural network learning capability that arises when synaptic weights are enabled to store details of the learning strategy. (**A**) The neural network is a generic recurrent network of spiking neurons, some of which exhibit SFA. The network was provided cue patterns and pattern deletion signals as input, and produced the appropriate completed pattern as outputs. (**B**) Demonstration of the capability of the network to learn 3 new attractors in one shot so that they can be used for input completion, and also to delete one of the attractors (here attractor 1) in one shot.

The recurrent network consisting of 300 spiking neurons, half of which exhibited SFA, was trained in the outer loop of L2L to be able to memorize any arbitrary three prototype patterns instantaneously in phase A. The patterns were randomly generated 25-bit patterns, and network performance was evaluated for patterns that did not occur during training in the outer loop. Then in phase B it could use these stored prototype patterns for completing partial network inputs. The network also was able to delete any of the three pattern prototypes (here pattern 1) in phase C, and to continue pattern completion with the remaining two pattern prototypes 2 and 3 (phase D). Note that the same partial network input or cue that lead in phase B to pattern (attractor) 1 is now completed in phase D to the next best prototype pattern 3 (with closest hamming distance).

The network was able to perform this four phase task for arbitrary prototype patterns consisting of 25 bits, achieving for new patterns a bitwise completion accuracy of 97.34% in phase B and 77.52% in phase D (for an average of 87.43% in both phases). See Methods for full details.

### 3.3 Fast adaptation of motor predictions

The brain is able to adapt its motor control commands very fast, sometimes even in a single trial (Crevecoeur et al., 2020a,b). It is unlikely that synaptic plasticity can accomplish that (Perich et al., 2018), and an alternative model has been missing. We show that one-shot adaptation of motor prediction can be achieved if synaptic plasticity in the outer loop of L2L is complemented by the capability to transiently store salient information in the network state. We demonstrate this for the case of a forward model for an arm. The brain needs such forward models to plan movements, and also to take corrective actions if needed (Lalazar and Vaadia, 2008; Wolpert and Ghahramani, 2000). Visual and proprioceptive feedback provide essential feedback for that (Wong et al., 2012).

We address the question of how the brain can quickly adapt its motor predictions for arm movements when kinematic or dynamic properties of the arm change. For example, carrying a load changes the distribution of masses over the arm, and using a tool in the hand changes its effective length. And yet, neural networks of the brain can quickly correct for these changes — without requiring multiple rounds of trial and error. Our goal was to produce a model of how RSNNs can achieve this without synaptic plasticity.

Here, we consider the case of a two-link arm as illustrated in Fig. 3B. The tip of the arm is moved by applying torques to each of the two joints. Both of its limbs are also subject to gravity. The task was to predict the angles of the two joints. But the masses and lengths of the two limbs were different in every episode, leading to very different trajectories even when the same torques were applied (Fig. 3C). The RSNN received as input the control torques applied to the arm model encoded through the population activity of 100 spiking neurons (see Fig. 3D and Methods). No direct information about the masses and lengths of the limbs were provided to the model, only the true angles of the limbs were given as feedback to the network with a delay of 100 ms (this feedback was set to 0 for the first 100 ms). Nevertheless the RSNN was able to adapt its predictions for a new arm within about 600 ms while moving it for the first time (Fig. 3E, F). This is substantially faster than previous models for adaptation of a forward model based on synaptic plasticity (Gilra and Gerstner, 2017). Essential for this fast adaptation was that the RSNN model included neurons with SFA, and that its synaptic weights were trained in the outer loop of L2L to enable this very fast adaptation (see Methods for details).

**Figure 3:**
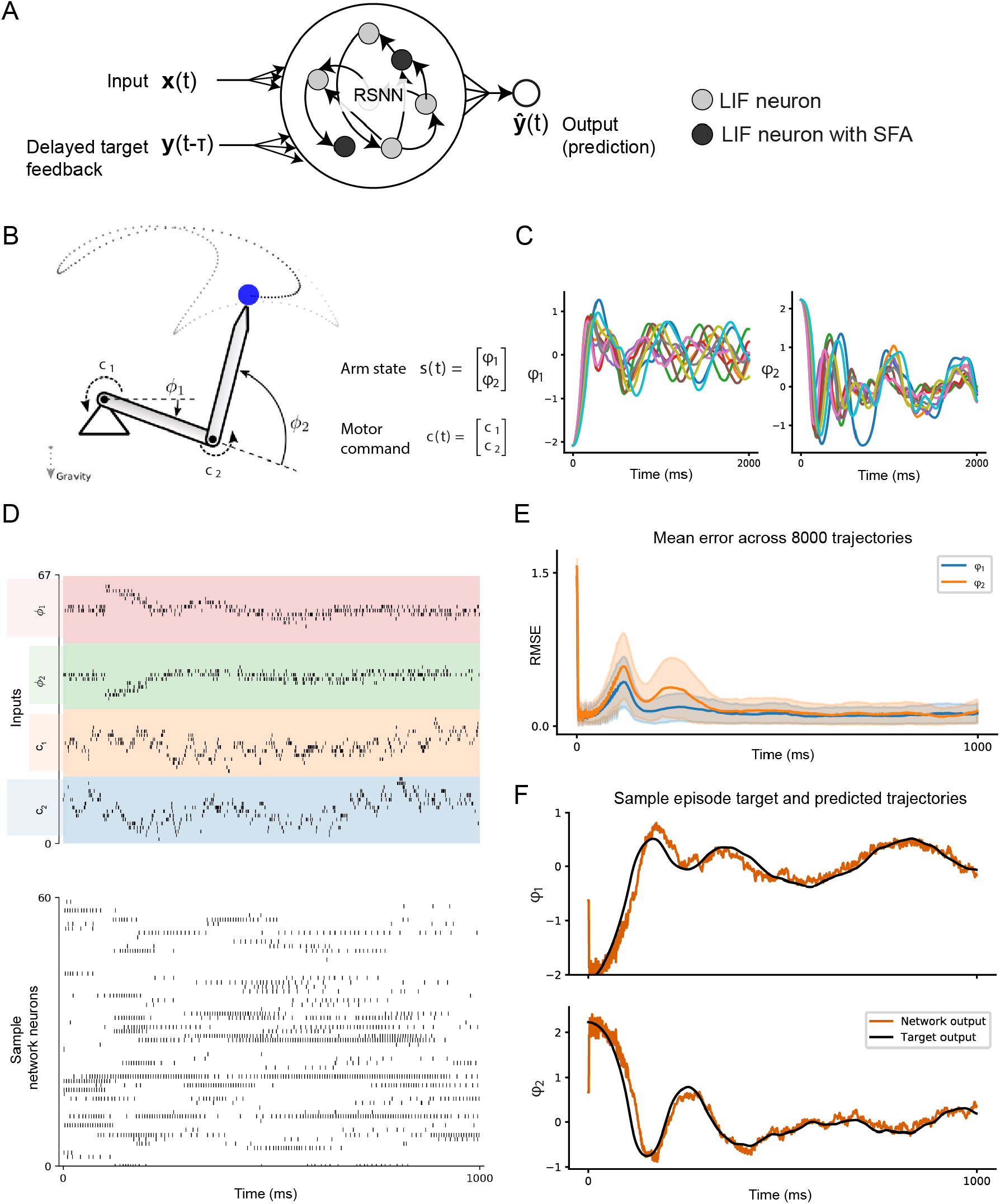
One-shot adaptation of a forward model for an arm. (**A**) The network setup for this and Sec. 3.4 consists of a generic recurrent network of spiking neurons some of which exhibit SFA. The network receives, in addition to the input *x*(*t*), a delayed target feedback signal *y*(*t* − *τ*), and produces the output *ŷ*(*t*) (where *y*(*t*) is the actual target). (**B**) Illustration of the two-link arm model with states given by the angle of the links, and the motor command applied on both the joints. (**C**) Sample trajectories generated for the same torque sequence by arms with different masses and lengths of the limbs. (**D**) Sample spike raster of neurons in the RSNN during an episode. The inputs to the network are, from top to bottom, the *τ* =100 ms delayed states *ϕ*_1_(*t* − *τ*) and *ϕ*_2_(*t* − *τ*) given as feedback; and the motor commands *c*_1_(*t*) and *c*_2_(*t*). (**E**) Root mean-squared error over all test episodes during the 1 second of inner-loop learning. (**F**) Target trajectories and network prediction for one sample test episode for an arm with new link lengths and masses.

### 3.4 Priors encoded in synaptic weights can significantly speed up learning

To demonstrate that synaptic weights can also be used to encode innate or previously learnt priors that can enable and/or speed up learning of complex tasks (point 2 of the Introduction), we will use a simple task where the RSNN has to learn a mapping *f* from input values *x* to output values *y* from example pairs (*x, y*) with *y* = *f* (*x*). Here, each task *C* corresponds to a mapping *f*. This learning task requires generalization from mappings *f* ^*/*^ that occurred during training to mappings *f* that did not occur during training. Obviously, this generalization is impossible if the learner has no prior knowledge about the function *f* that is to be learnt. Artificial neural networks with continuous activation functions implicitly use a prior that the target function *f* is smooth. But SNNs do not automatically apply a smoothness prior, since they can just as well represent discontinuous input-output mappings. We wondered whether the weights of a RSNN could encode a smoothness prior, and possibly further structural properties of potential target functions *f*.

We focused on the specific case where it is a priori known that the target function *f* is a sinus function, but with unknown phase and amplitude. In each inner loop episode, the RSNN received a sequence of inputs *x*^*k*^ from some mapping *f*, each encoded through the population activity of 100 spiking neurons for 20 ms. In addition to *x*^*k*^, it received the target output *y*^*k−*1^ = *f* (*x*^*k−*1^) for the previously presented input (see Fig. 1B and Methods). In this way, the network received a delayed feedback about its desired output which it could use to adapt its behavior in accordance with its internal prior on the family of functions *f*. The network had to predict the target *y*^*k*^ = *f* (*x*^*k*^) at each time step *k*, and was trained in the outer loop to do so in batches of episodes with a different mappings/tasks. During testing, previously unseen mappings were used.

Fig. 4B-E demonstrates that this prior information can in fact be encoded in the synaptic weights of the RSNN, which are determined in the outer loop of L2L. We applied a simple trick for visualization to make the prior or internal model of the RSNN at any moment in time visible: see the orange lines in Fig. 4B-E. These orange lines show for any potential input value *x* (in the domain of *f*) the output *y* which the RSNN would give if this *x* would occur as the next network input (in a hypothetical experiment, that has no effect on the next steps of the (inner loop) learning process for learning the target function *f*). More precisely, the network state is not allowed to change when these test inputs *x* are shown.

**Figure 4:**
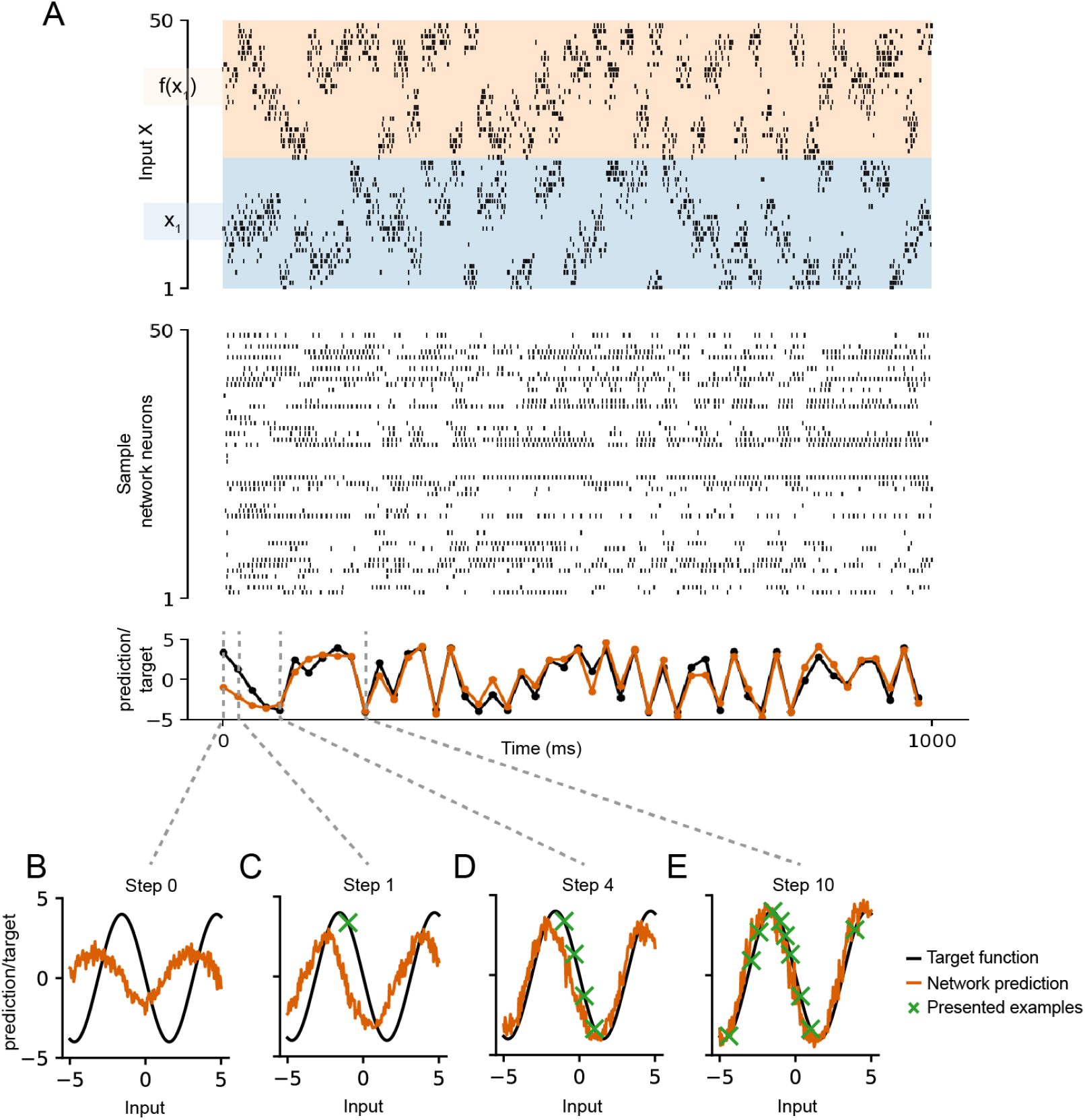
Encoding structural priors for learning. (**A**) Sample spike raster of neurons in the RSNN during an episode of learning to predict a previously unseen sinusoidal curve. (**B-E**) Snapshots of the internal model of the network at different inner loop steps, illustration of the prior knowledge acquired by the RSNN containing LIF neurons with SFA through L2L for the family ℱ of all sinus functions with different phases and amplitudes, but a fixed frequency. Orange curves show the effective internal model of the RSNN at different stages of learning from examples of the target function (marked by green crosses).

Fig. 4B shows the prior or internal model that is engraved into the RSNN through its synaptic weights before it has received any training example ⟨*x, y*⟩. One clearly sees that this internal model is in fact a sinus function. In addition, this internal model already reflects the frequency of the target sinus function *f*, since this is the same for all potential target functions that were considered in the outer loop of L2L. The subsequent panels, Fig. 4C-E, show that the internal model is updated when some actual training examples — indicated by green crosses — are received by the RSNN. One sees from Fig. 4C that a single training example brings the internal model already quite close to the function *f* from which the training examples are generated. Fig. 4D shows that the prior of the RSNN has such a strong impact that even when it receives 4 training examples from *f* that happen to lie approximately on a straight line, its internal model (i.e, posterior) still has the form of a sinus function, rather than a straight line. Fig. 4E shows the internal model once the network has received sufficient number of points to fully predict the sinus function. The normal temporal progression of the experiment at each step is shown in Fig. 4A, where the network state progresses normally after each example (that were marked by green crosses in Fig. 4B-E) is presented to the network.

### 3.5 Spiking neural networks can learn extremely fast from rewards — without engaging synaptic plasticity

We now demonstrate the ability of synaptic weights to encode innate or previously learnt priors that can enable and/or speed up learning of complex reinforcement learning tasks. For this, we use variations of the well-known Morris water-maze task (Morris, 1984; Vasilaki et al., 2009) to define the range ℱ of learning tasks for L2L. Here the subject has to learn to find a target in a 2D arena, and to navigate subsequently to this target from random positions in the arena (Fig. 5A, B).

**Figure 5:**
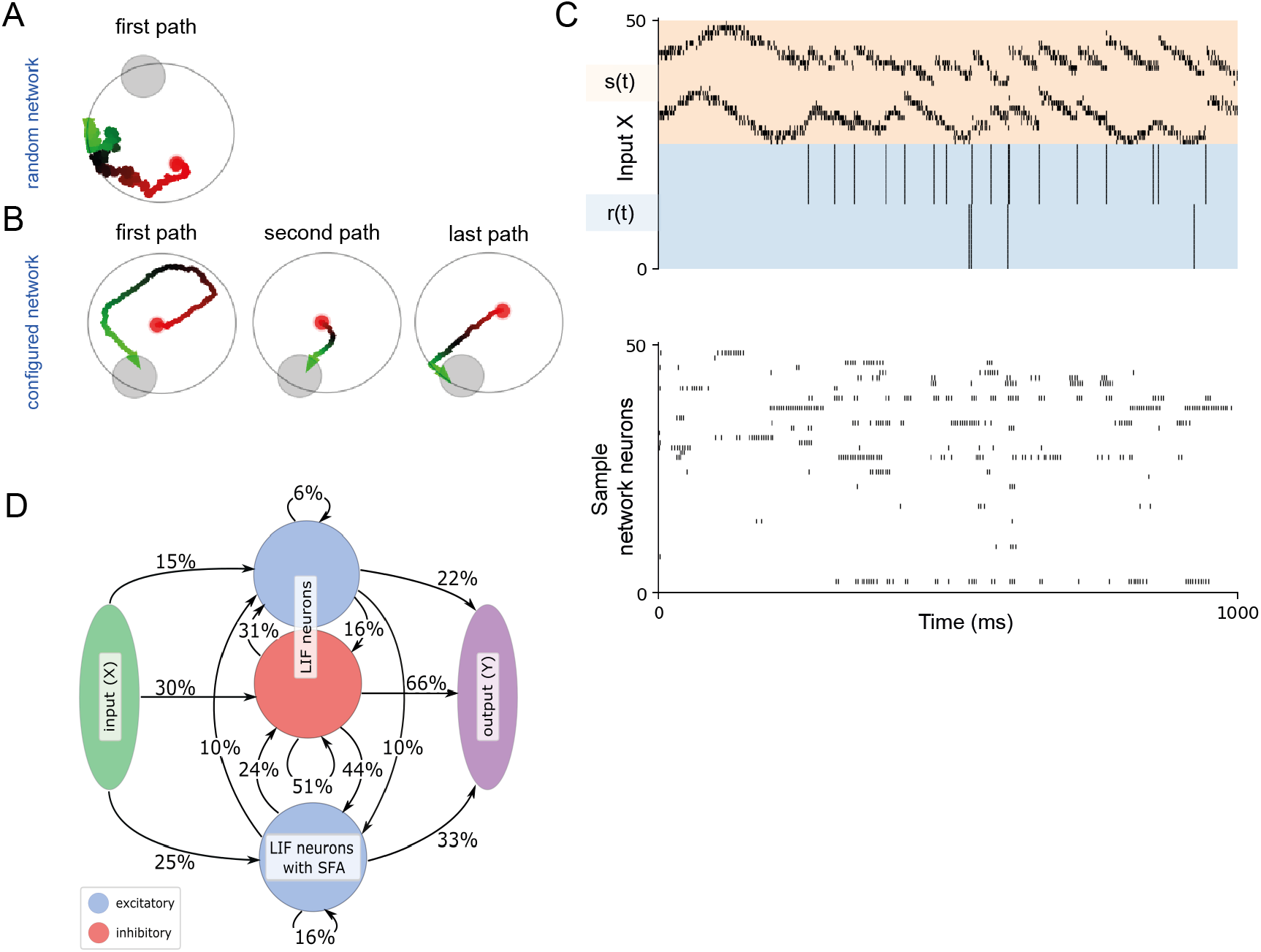
Reward-based learning of an RSNN without synaptic plasticity. (**A, B**) Samples of navigation paths produced by the RSNN before and after this training. Before training, the agent performs a random walk (**A**). In this example it does not find the goal within the limited episode duration. After tuning the synaptic weights of the RSNN in the outer loop of L2L (**B**), the RSNN had acquired an efficient exploration strategy that uses two pieces of abstract knowledge: that the goal always lies on the border, and that the goal position remained the same throughout an episode. Note that all synaptic weights of the RSNNs remained fixed during an episode. (**C**) The network dynamics that produced the behavior shown in (**B**). (**D**) The architecture of the RSNN, consisting of excitatory and inhibitory neurons, with 20% connectivity obeying Dale’s law. Synaptic connectivity was optimized within these constraints in the outer loop of L2L.

The task consists of episodes that each last 2 seconds. The goal was placed randomly for each episode on the border of the arena. When the agent reached the goal, it received a reward of +1, and was placed back randomly in the arena. When the agent hit a wall, it received a negative reward of −0.02 and the velocity vector was truncated to remain inside the arena. The objective was to maximize the number of goal reaches within an episode. The Morris water-maze task is related to one of the more challenging demos of (Wang et al., 2016) and (Duan et al., 2016) in applying L2L to networks of LSTM units. But it had remained open whether this learning paradigm can also be applied to biologically more realistic neural network models. We investigate here to what extent a sparsely connected network of excitatory and inhibitory neurons that observes Dale’ law can learn to solve the Morris water-maze task within a few trials.

We are addressing here at the same time point 2 of the Introduction: Can synaptic weights encode common structure in this family of task so that the network can use this abstract knowledge for more efficient learning? Concretely, there are two pieces of abstract knowledge in the family of water-maze tasks: The fact that goals are always on the border of the arena, and the fact that the goal position is constant within each episode. Note that we did not allow synaptic plasticity to take place during the short duration of a testing episode, only in the outer loop of L2L.

Since RSNNs with just a few hundred neurons are not able to process visual input, we provided the current position of the agent within the arena through a place-cell like Gaussian population rate encoding of the current position (see orange segment in the top row of Fig. 5C). Note that the lack of visual input already makes it challenging to move along a smooth path, or to stay within a safe distance from the wall. The agent received information about positive and negative rewards in the form of spikes from external neurons (blue segment of the upper row of Fig. 5C). We used the outer loop of L2L to configure the network to solve this task as fast as possible — see Methods for details of the optimization process used to configure the network. In this task the RSNN had 400 recurrent units (200 excitatory and 80 inhibitory standard LIF neurons, and 120 excitatory neurons with SFA) and a synaptic connectivity of 20%. We allowed the network to rewire itself during the outer loop of L2L, which substantially improved the performance. The resulting network diagram and spike raster is shown in Fig. 5D. A performance comparison with a random network with the same overall number of synaptic connections is given in Supplementary Fig. 1-1.

The first path in Fig. 5B shows that the RSNN is able to make use of the fact that the goal is located on the border of the maze. The second and last paths show that the RSNN also makes use of the abstract knowledge that the goal position remains fixed during an episode. Fig. 5C exhibits sample spike trains from excitatory and inhibitory LIF neurons without SFA, and of excitatory LIF neurons with SFA.

Altogether this demo shows that RSNN are able to learn very fast from rewards, without engaging synaptic plasticity. Furthermore it shows that synaptic weights of SNNs can encode abstract knowledge which makes learning of a behaviour substantially more efficient.

## 4 Discussion

We have revisited the roles of synaptic plasticity and network dynamics for learning in spike-based models of recurrent neural networks in the brain. So far most biologically plausible models for learning in RSNNs have focused on STDP or other synaptic plasticity mechanisms. Usually it was also assumed that this synaptic plasticity mechanisms becomes immediately effective, which is not consistent with experimental data on STDP (Froemke et al., 2010). Our results suggest that such mechanisms for synaptic plasticity are likely to be complemented with other mechanisms that especially support fast learning.

One fundamental insight that emerges from this analysis is that learning in RSNNs can be substantially more versatile and faster than previously thought. In particular, salient information during learning can also be encoded in the hidden variables of neurons if one also includes slower processes of biological neurons in the neuron models. We have considered here only one such slow process, spike frequency adaptation of neurons, and shown that it has a remarkable impact on the learning capability of a network of spiking neurons. In particular one arrives in this way at the first spiking neural network models that can explain, through a biologically realistic neural network model, the capability of brains to learn significant behavioural improvements in very few trials, often even in a single trial. Our neural network model is based on data-based models for neurons, such as the GLIF neurons of (Billeh et al., 2020). Hence we conjecture that our new learning paradigms can also be implemented and tested in such large-scale data-based models of brain areas. It particular, it opens the door for modelling concrete fast learning processes of the brain that have been studied for example in (Perich et al., 2018) and (Crevecoeur et al., 2020a). Our model can be used to understand these and related biological phenomena, such as fast adaptation of motor predictions, see Fig. 3.

We have demonstrated two specific advantages of this new model for learning in recurrent networks of spiking neurons:

1. It substantially enlarges the diversity and power of learning strategies by which recurrent networks of spiking neurons can learn, see for example the demonstration with one shot learning of attractors by a RSNN and instantaneous deletion of an attractor in Fig. 2.
2. These networks can learn substantially faster than previously thought by making use of prior knowledge that is stored in their synaptic weights, see Fig. 4 and 5. In particular, we have shown in Fig. 4 that, once the network has learnt the overall task structure, it is able to ignore misleading information, thereby enabling robust learning from few examples (Gershman and Niv, 2010).

We have also shown in Fig. 5 that an application of our two-tiered learning model can solve the Morris water maze task, a well known biological learning paradigm (Morris, 1984; Vasilaki et al., 2009). This task was modelled as a continuous control problem, and we applied meta-reinforcement learning to the spiking neural network. This enabled the outer loop to extract two abstract pieces of information into the spiking neural network from its interaction with the environment: that the goals are always on the perimeter of the maze, and that the goal position does not change during trials that belong to the same learning episode. The network was able to apply this abstract learnt knowledge in order to solve very fast instances of the task that it had never encountered before.

Our new model for learning in neural networks of the brain makes a clear experimentally testable predictions: It predicts that traces of fast learning become first apparent in a modified network dynamics, and only later in modifications of synaptic strengths. More specifically, our model predicts that very recently acquired information can be decoded first from the effective firing thresholds of neurons or other slowly changing variables of neurons and synapses. We expect that some of this newly acquired information is transformed and generalized during consolidation into synaptic weights (Stickgold and Walker, 2013).

Altogether our results suggest that learning in RSNNs of the brain is likely to engage other neurophysiological mechanisms besides synaptic plasticity, and that evolution, development and prior learning are likely to have configured and aligned these different processes so that they complement each other when a new learning task arises. This perspective opens the door to a much richer and functionally more powerful range of network learning methods than those which just consider synaptic plasticity. The spiking neural networks that emerged in the various tasks we considered, computed and learnt with a brain-like sparse firing activity. This is quite different from the dynamics of a spiking neural networks that operates with rate-codes. Hence these paradigms also broaden our insight into ways in which brains are able to compute and learn with sparsely active spiking neurons.

## Acknowledgements

This research was partially supported by the Human Brain Project (Grant Agreement number 785907) of the European Union and by the ERA-NET CHIST-ERA programme by the Austrian Science Fund (FWF) (project SMALL, project number I 4670-N). We gratefully acknowledge the support of NVIDIA Corporation with the donation of the Quadro P6000 GPU used for this research. Computations were carried out on the Human Brain Project PCP Pilot Systems at the Juelich Supercomputing Centre, which received co-funding from the European Union (Grant Agreement number 604102) and on the Vienna Scientific Cluster (VSC).

