## Supplemental Figure 5-1 for "Revisiting the role of synaptic plasticity and network dynamics for fast learning in spiking neural networks"

### Extended data for: Revisiting the role of synaptic plasticity and network dynamics for fast learning in spike-based neural network models

Anand Subramoney[1,2], Guillaume Bellec[1,3], Franz Scherr[1], Robert Legenstein[1], and Wolfgang Maass[1,*]

[1]Institute for Theoretical Computer Science, Graz University of Technology, Austria
[2]Institute for Neural Computation, Ruhr University Bochum, Germany
[3]Laboratory of Computational Neuroscience, Ecole Polytechnique Fédérale de Lausanne (EPFL), Switzerland

January 25, 2021

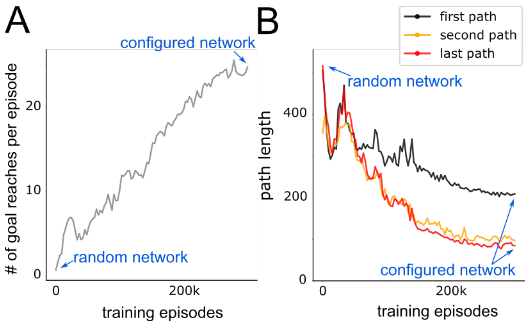


Figure 5-1: **Reward-based learning of an SNN without synaptic plasticity.** (**A, B**) The performance of the network is shown as the training in the outer loop progresses, starting from a completely random network at training episode 0 to the fully configured network at the final training episode. Initially the network is unable to reach the goal more than a few times in an episode. But as the outer loop training progresses, the network is able to reach the goal more often per episode. Moreover, a clear difference between the first path and the second to last path is visible in (**B**) at the end of training: the agent explores the arena to find the goal in the first path, but from the second path onwards, the agent goes directly to the goal, exploiting the knowledge of the position of the goal acquired in the first path.
